# Sequence Comparison without Alignment: The *SpaM* approaches

**DOI:** 10.1101/2019.12.16.878314

**Authors:** Burkhard Morgenstern

**Affiliations:** University of Goettingen

## Abstract

Sequence alignment is at the heart of DNA and protein sequence analysis. For the data volumes that are nowadays produced by massively parallel sequencing technologies, however, pairwise and multiple alignment methods have become too slow for many data-analysis tasks. Therefore, fast alignment-free approaches to sequence comparison have become popular in recent years. Most of these approaches are based on *word frequencies*, for words of a fixed length, or on word-*matching* statistics. Other approaches are based on the length of *maximal word matches*. While these methods are very fast, most of them are based on ad-hoc measures of sequences similarity or dissimilarity that are often hard to interpret. In this review article, I describe a number of alignment-free methods that we developed in recent years. Our approaches are based on *spaced word matches (‘SpaM’)*, i.e. on inexact word matches, that are allowed to contain mismatches at certain pre-defined positions. Unlike most previous alignment-free approaches, our approaches are able to accurately estimate phylogenetic distances between DNA or protein sequences based on stochastic models of molecular evolution.

## 1 Introduction

Alignment-free sequence comparison has a long tradition in bioinformatics. The first approaches to compare sequences without alignments were proposed in the Nineteen-eighties by E. Blaisdell [6, 7]. The interest in alignment-free methods increased when more and more partially or completely sequenced genomes became available through novel sequencing technologies, leading to an urgent need for faster methods of sequence comparison. Most existing alignment-free methods represent sequences as *word-frequency vectors* for words of a fixed length *k* so-called *k*-mers –, and by comparing *k*-mer frequencies instead of comparing sequences position-by-position, based on alignments [75, 32, 70, 13, 79]. This approach has been extended by taking the background word-match frequencies into account [62, 80, 73, 1]; a review of these latter methods is given in [63]. Other approaches to alignment-free sequence comparison are based on the length of *maximal common sub-words* of the compared sequences, to define alternative measures of sequence similarity or dissimilarity [77, 14, 45, 61, 76].

The main advantage of these word-based methods is their high speed, compared to alignment-based methods. While – for most scoring schemes – finding an optimal alignment of two sequences takes time proportional to the *product* of their lengths [56, 22, 51], word-based or alignment-free methods are much more efficient, since word-frequency vectors can be calculated in time proportional to the length of the analyzed sequences. Similarly, the length of longest common sub-words can be efficiently found using data structures such as *generalized suffix trees* or *suffix arrays* [25]. A review of earlier alignment-free methods is given in [78]; more recent review papers are [27, 72, 84, 5, 40]. A first systematic benchmark study of alignment-free methods has been published in 2019, as a collaboration of several groups working in the field [83].

From the beginning, *phylogenetic tree reconstruction* has been a main application of alignment-free sequence comparison. Choi and Kim, for example, were able to calculate a phylogenetic tree of > 4, 000 whole-proteome sequences [12], using the alignment-free tool *FFP* that has been developed by the same group [70]. The fastest phylogeny methods are *distance-based* approaches: to calculate a tree for a set of taxa, these methods use pair-wise distance values as input. Thus, for each pair of compared taxa, their distance or dissimilarity needs to be measured in some way. Once all pairwise distances have been calculated for the input set of taxa, a matrix with these distances can be used as input for standard distance-based methods for phylogeny reconstruction. The most commonly used methods for distance-based phylogeny reconstruction are *Neighbor-Joining (NJ)* [67] and *BIONJ* [24].

If DNA sequences are compared, a common way of defining the distance between two evolutionarily related sequences is to use the (estimated) number of *substitutions per position* that have occurred since the two sequences have evolved from their last common ancestor. The simplest substitution model for nucleic-acid sequences is the *Jukes-Cantor* model. Here, all nucleotide substitutions are assumed to occur with the same probability per time unit. Under this model, the number of *substitutions per position* can be estimated from the *number of mismatches per position* in an alignment of the compared sequences, using the well-known *Jukes-Cantor formula* [37]. More elaborate substitution models are available to for DNA or protein sequences, that account for different substitution probabilities for different pairs of nucleotide or amino-acid residues.

By contrast, most of the above mentioned alignment-free methods are not based on probabilistic models of evolution, they rather use heuristic measures of sequence similarity or dissimilarity. If sequences are represented by word-frequency vectors, for example, standard distance measures on vector spaces can be applied to these frequency vectors, in order to calculate a ‘distance’ between two compared sequences, such as the *Euclidean distance* or the *Manhattan distance*. Such distances, however, are hard to interpret from a phylogenetic point-of-view. Clearly, a pair of closely related sequences will have more words in common – so the Euclidean distance between their word-frequency vectors will be smaller – than is the case for a more distantly related pair of sequences.

But distance values calculated in this way are not measures of evolutionary distances, e.g. in terms of events that happened since two sequences separated from their last common ancestor. They only indicate if one pair of sequences shares more or less similarity than another pair of sequences. Such heuristic distance measures can be used for clustering, but not for more accurate phylogenetic analyses.

Since the distance values calculated by earlier word-based alignment-free methods have no direct phylogenetic interpretation, it would make no sense to ‘evaluate’ the accuracy of these values directly. Therefore, the developers of these methods did not evaluate and benchmark the distance values produced by their methods themselves. Instead, they applied clustering algorithms or distance-based tree reconstruction methods to these distances and evaluated the resulting trees. Again, since the computed distances between the sequences have no direct meaning, the branch-lengths of these trees were usually ignored, and only the resulting *tree topologies* were evaluated. The standard approach to evaluate tree topologies is to compare them to trusted reference *topologies* using the *Robinson-Foulds* metric [65]. Note that this is only a very rough and indirect method to benchmark the performance of sequence-comparison methods.

It was only in the last ten years or so that alignment-free methods were proposed that can estimate phylogenetic distances in the sense of an underlying probabilistic model of sequence evolution. The first such approach has been published in 2009 by Haubold et al. [29]. These authors developed *kr*, an alignment-free method that can accurately estimate phylogenetic distances between *DNA* sequences [29] in the sense of the *Jukes-Cantor* model. That is, *kr* estimates the number of nucleotide substitutions per sequence positions since the compared sequences have evolved from their last common ancestor. To this end, the program used the average length of common substrings between the compared sequences. Later, we proposed an approach to estimate phylogenetic distances based on the length distribution of *k*-mismatch common substrings [53].

In the last few years, other alignment-free methods have been proposed to estimate phylogenetic distances in a rigorous way [82, 28, 47, 39]. Some of these methods are based on so-called *micro-alignments*, short gap-free pairwise alignments with a simplistic structure, that can be rapidly calculated. So, strictly spoken, these methods are not *alignment-free*. They are referred to as ‘alignment-free’ anyway, since they avoid the time-consuming process of calculating optimal alignments over the entire length of the input sequences. Other approaches to estimate distances in a stochastically rigorous way are based on the number of word matches [55]. More recently, extremely fast programs have become popular that can accurately estimate phylogenetic distances between DNA sequences from the number of word matches, using the so-called *Jacard Index* [36] and *min-hash* algorithms [10]. A widely-used implementation of these ideas is *Mash* [58]; further improvements to this approach have been proposed and are implemented in the programs *Skmer* [68], *Dashings* [4] and *Mash Screen* [57].

## 2 Spaced Words

In 2013, we proposed to use so-called *spaced words* that contain *wildcard* characters at certain positions, for alignment-free *DNA* and protein sequence comparison [8, 43, 33]. A spaced word is based on a pre-defined binary pattern *P* that are called *match positions* (’1’) and *don’t-care positions* (’0’). Given such a pattern *P*, we defined a spaced word *w* with respect to *P* to be a word that has the same length as the pattern *P* and that has symbols for nucleotide or amino-acid residues at the *match positions* of *P* and *wildcard symbols* (‘*’) at the *don’t-care positions*, see Figure 1 for an example. Spaced words – or *spaced seeds* – have been previously introduced in database searching, to improve the sensitivity of the standard *seed-and-extend* search strategy [49]. Efficient algorithms have been proposed to optimize the underlying patterns [35], and for *spaced-seed hashing* [60].

**Figure 1:**
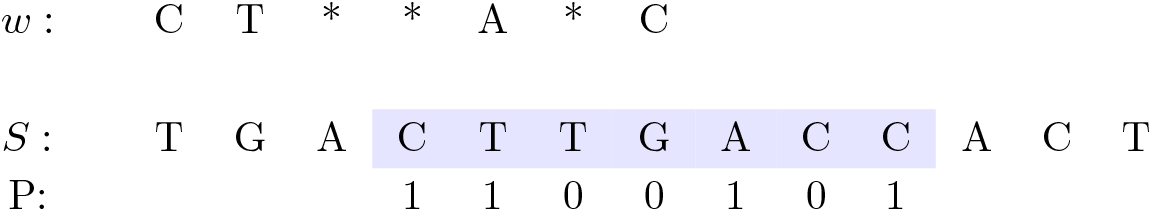
Spaced word *w* with respect to a pattern *P* = 1100101 of length *ℓ* = 7. *w* consists of nucleotide symbols at the *match positions* (’1’) of *P* and of wildcard symbols, represented as ′*′ at the *don’t-care positions* (’0’). *w* occurs at position 4 in the *DNA* sequence *S*.

In a first study, we simply replaced *word-frequency* vectors by *spaced-word* frequency vectors to calculate distances between DNA and protein sequences. As in earlier word-based methods, we used the *Euclidean* distance or, alternatively, the *Jenson-Shannon* distance of the frequency vectors to define the distance between two DNA or protein sequences. As a result, the quality the resulting phylogenetic trees was improved, compared to when we used contiguous words – in particular when we used *multiple* binary patterns and the corresponding spaced-word frequencies, instead of single pattern [43]. The resulting software program is called *Spaced Words*, or *Spaced*, for short.

Our spaced-words approach was motivated by the *spaced-seeds* [48] that have been introduced in database searching, to improve the sensitivity of *hit-and-extend* approaches such as *BLAST* [2]. The main advantage of spaced word matches – or ‘spaced seeds’ – compared to contiguous word matches is that neighbouring spaced-word matches are statistically less dependent, so they are distributed more evenly over the sequences. In database searching, this increases the *sensitivity*, i.e. the probability of finding sequence similarities. For alignment-free sequence comparison, we have shown that results obtained with spaced words are statistically more stable than results based on contiguous words [55].

Note that, like earlier alignment-free approaches, this first version of the program *Spaced* was still based on heuristic measures of sequence similarity. It does not estimate evolutionary distances in the sense of some probabilistic model. Later, however, we introduced a new distance measure in *Spaced* based on the number of spaced-word matches [55] that actually estimates phylogenetic distances between DNA sequences in the sense of the *Jukes-Cantor* model [37]. This is now the the default distance measure used in the program *Spaced*. To find good patterns – or *sets* of patterns in the *multiple-pattern* approach –, we developed a program called *rasbhari* [26].

## 3 *Filtered Spaced-Word Matches* and *Prot-SpaM*

In a subsequent project, we introduced a different approach to use spaced words for alignment-free sequence comparison. Instead of comparing spaced-word *frequencies*, we used *spaced-word matches (SpaM)* as a special type of *micro-alignments*. For a binary pattern *P* as above, a *SpaM* between two sequences is simply the occurrence of the same spaced word in both sequences with respect to *P*, see Fig. 2 for an example. In other words, a *SpaM* is a local, gap-free alignment that has the same length as the pattern *P* and that has matching nucleotides or amino acids at the *match positions* and possible mismatches at the *don’t-care positions* of *P*. The idea is to consider a large number of *SpaMs*, and to estimate phylogenetic distances between two sequences by looking at the residues that are aligned to each other at the *don’t-care positions* of these *SpaMs*. Obviously, this is only possible if the considered *SpaMs* represent true homologies, so we have to filter out spurious background *SpaMs*. To do so, our program first considers all *possible SpaMs* between two input sequences and calculate a score for each *SpaM* based on the aligned residues at the *don’t-care* positions. The program then discards all *SpaMs* with scores below some threshold. We could show that, that with this sort of *SpaM filter*, one can reliably separate true homologies (’signal’) from random *SpaM* (’noise’).

**Figure 2:**
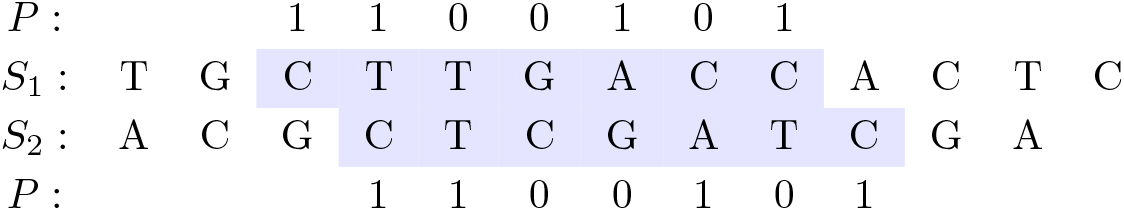
*Spaced-word match (SpaM)* between two DNA sequences *S*_1_ and *S*_2_ with respect to a binary pattern *P* = 1100101 of length *ℓ* = 7, representing *match positions* (‘1’) and *don’t-care positions* (‘0’). The two segments have matching nucleotides at all *match positions* of *P* but may mismatch at the *don’t-care* positions.

An implementation of this approach for *DNA* sequences is called *Filtered Spaced Word Matches (FSWM)*, To estimate distances between *DNA* sequences, *FSWM* calculates the proportion of mismatches at the *don’t-care* positions of the selected *SpaMs*, as an estimate of the proportion of mismatches in the (unknown) full alignment of the two sequences. It then applies the usual *Jukes-Cantor* correction, to calculate the estimated *number of substitutions per position* since the two sequences have evolved from their last common ancestor. By default, the program uses a pattern *P* of length *ℓ* = 112 with 12 *match positions* and 100 *don’t-care* positions, but the user can adjust these parameters. The length of the pattern *P* seems to be certain limitation, as it means that, by default, the program is restricted to using gap-free homologies of length ≥ *ℓ* = 112. A sufficient number of *don’t-care* positions is necessary, though, to reliably distinguish *SpaMs* representing true homologies from random background *SpaMs*. To speed-up the program, it can be run with multiple threads; by default 10 threads are used.

An implementation of the same algorithm for protein sequences is called *Prot-SpaM* [46]. Here, we are using by default a pattern with 6 *match positions* and 40 *don’t-care positions*, i.e. with a length of *ℓ* = 46. For protein sequences, we are using the *BLOSUM 64* substitution matrix [30] to score *SpaMs* in order to filter out low-scoring random *SpaMs*. Again, the user can modify these parameters. To estimate the evolutionary distance between two protein sequences, we consider the pairs of amino acids aligned to each other at the don’t-care positions of the selected spaced-word matches, and we are using the *Kimura* model [38] that approximates the *PAM* distance [17] between sequences based on the number of mismatches per position.

## 4 *Read-SpaM:* estimating phylogenetic distances based on unassembled sequencing reads

Several authors have pointed out that alignment-free approaches can be applied, in principle, not only to full genome sequences, but also to unassembled reads. Some approaches have been particularly designed for this purpose [82, 68]. The ability to estimate phylogenetic distances based on unassembled reads is not only useful in phylogeny studies, but also in biodiversity research [68] or in clinical studies [20, 9]. Here, species or strains of bacteria can often be identified by *genome skimming*, i.e. by low-coverage sequencing [81, 21, 64, 19, 50, 68].

We adapted our *FSWM* approach to estimate phylogenetic distances between different taxa using unassembled reads; we called this approach *Read-SpaM* [42]. This software can estimate distances between an assembled genomes from one taxon and a set of unassembled reads from another taxon, or sets of unassembled reads from two different taxa. Using simulated sequence data, we could show that *Read-SpaM* can accurately estimate distances between genomes up to 0.8 substitutions per position, for sequence coverage as low as 2^−9^*X*, if an assembled genome is compared to unassembled reads. If unassembled reads from two different taxa are compared, distances estimates by *Read-SpaM* are still accurately for up to 0.7 substitutions per position, for a sequencing coverage down to 2^−4^*X*.

## 5 The most recent approaches: *Multi-SpaM* and *Slope-SpaM*

For nucleic-acid sequences, we extended our *Filtered-Spaced Words Matches* approach from pairwise to multiple sequence comparison [18]. Our software *Multi-SpaM* is based on spaced-word matches between four sequences each. Such a multiple spaced word match is, thus, a local gap-free *four-way alignment* with columns of identical nucleotides at the *match positions* of the underlying binary pattern *P*, while mismatches are, again, allowed at the *don’t-care positions* of *P*. An example is given in Fig. 3, such local four-way alignments are also called *P-blocks*.

**Figure 3:**
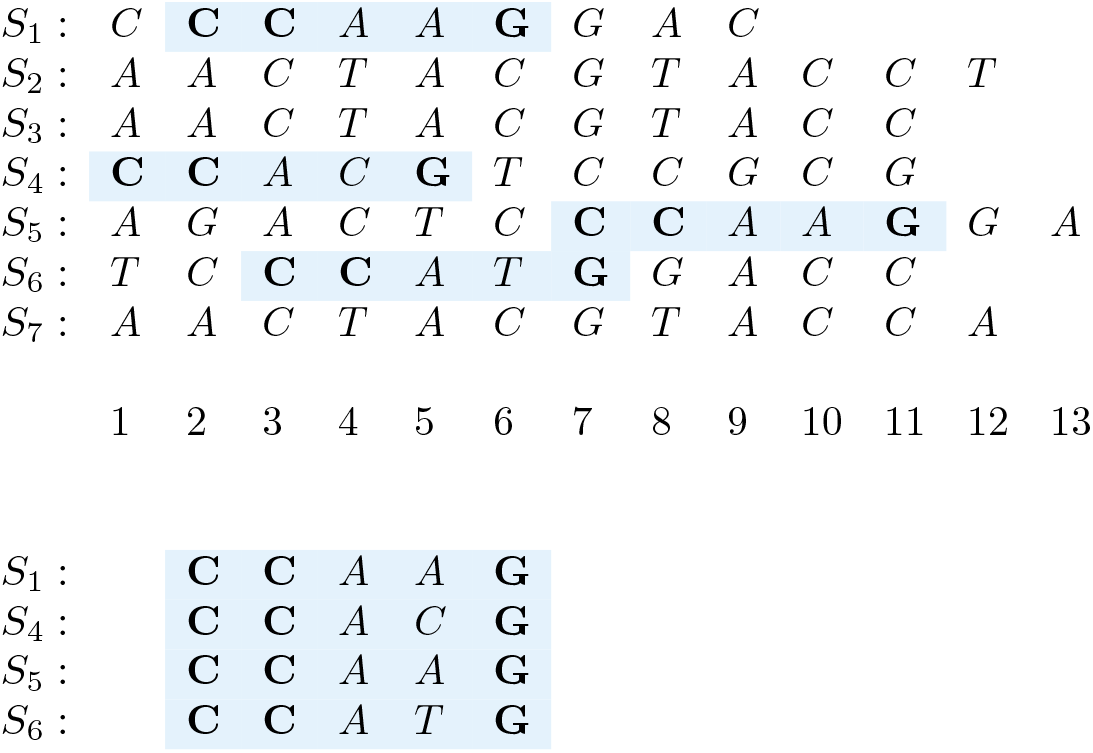
*P*-block for a pattern *P* = 11001: the spaced word *W* = *CC* ∗ ∗*G* occurs at [*S*_1_, 2], [*S*_4_, 1], [*S*_5_, 7] and [*S*_6_, 3] (top). A *P*-block defines a gap-free local four-way alignment (bottom).

*Multi-SpaM* samples *P*-blocks from the set of input sequences; by default up to 10^6^ *P*-blocks are sampled. To ensure that these *P*-blocks represent true homologies, only those *P*-blocks are considered that have a score above a certain threshold. For each of the sampled *P*-blocks, the program then uses *RAxML*[74] to calculate an optimal unrooted quartet tree topology. Finally, the program *Quartet MaxCut*[71] is used to calculate a super tree topology from these quartet topologies.

As another approach to alignment-free phylogeny reconstruction, we developed a program called *Slope-SpaM* [66]. This program considers the number *N_k_* of *k*-mer matches or *spaced-word* matches for patterns *P* with *k match positions*, respectively, between nucleic-acid sequences. It calculates *N_k_* for different values of *k* and estimates the *Jukes-Cantor* distances between sequences – i.e. the average number of substitutions per sequence position since the sequences diverged from their most recent common ancestor – from the decay of *N_k_* when *k* increases. This way, evolutionary distances can be calculated accurately for up to around 0.5 substitutions per sequence position.

## 6 Back to Multiple Sequence Alignment

There is not strict separation between sequence alignment and word-based, alignment-free methods. As mentioned above, a whole class of so-called ‘alignment-free’ methods are based on ‘micro-alignments’, local pairwise alignments of a simple structure, that can be rapidly calculated. In *Multi-SpaM*, we extended this approach to local *multiple* alignments.

Ironically, one of the first major applications of fast alignment-free methods was *multiple sequence alignment (MSA)*. The programs *MUSCLE* [23] or *Clustal Omega* [69], for example, are using word-frequency vectors to rapidly calculate *guide trees* for the ‘progressive’ approach to MSA [23]. Similarly, fast alignment-free methods are used to find *anchor points* [54, 34] to make alignments of large genomic sequences possible [31, 52, 41, 15, 16, 3, 59]. In a recent study [44], we used our program *FSWM* to generate anchor points for multiple genome alignment. We could show that, if distantly related genomes are compared, spaced-word matches are more sensitive and lead to better output alignments than anchor points that are based on exact word matches.

## Software Availability

We made *Filtered Spaced Word Matches (FSWM)* available through a web interface at *Göttingen Bioinformatics Compute Server (GOBICS)* at http://fswm.gobics.de/ see Figure 4. There are certain limitations at this web server for the size of the input data: (*a*) the upper limit for the total size of the input sequences is 512 *mb*, (*b*) the number of input sequences must be between 2 and 100, and the minimum length of each input sequence is 1000 *bp*. At our web server, the underlying pattern *P* has by default 12 *match positions* and 100 *don’t care positions*. The number of *match positions* can be adapted by the user. To calculate a *score* for each spaced-word match, a nucleotide substitution matrix published by Chiaromonte et al. [11] is used. By default, the cut-off value to distinguish ‘homologous’ from background spaced-word matches is set to 0. This value, too, can be adjusted by the user.

**Figure 4:**
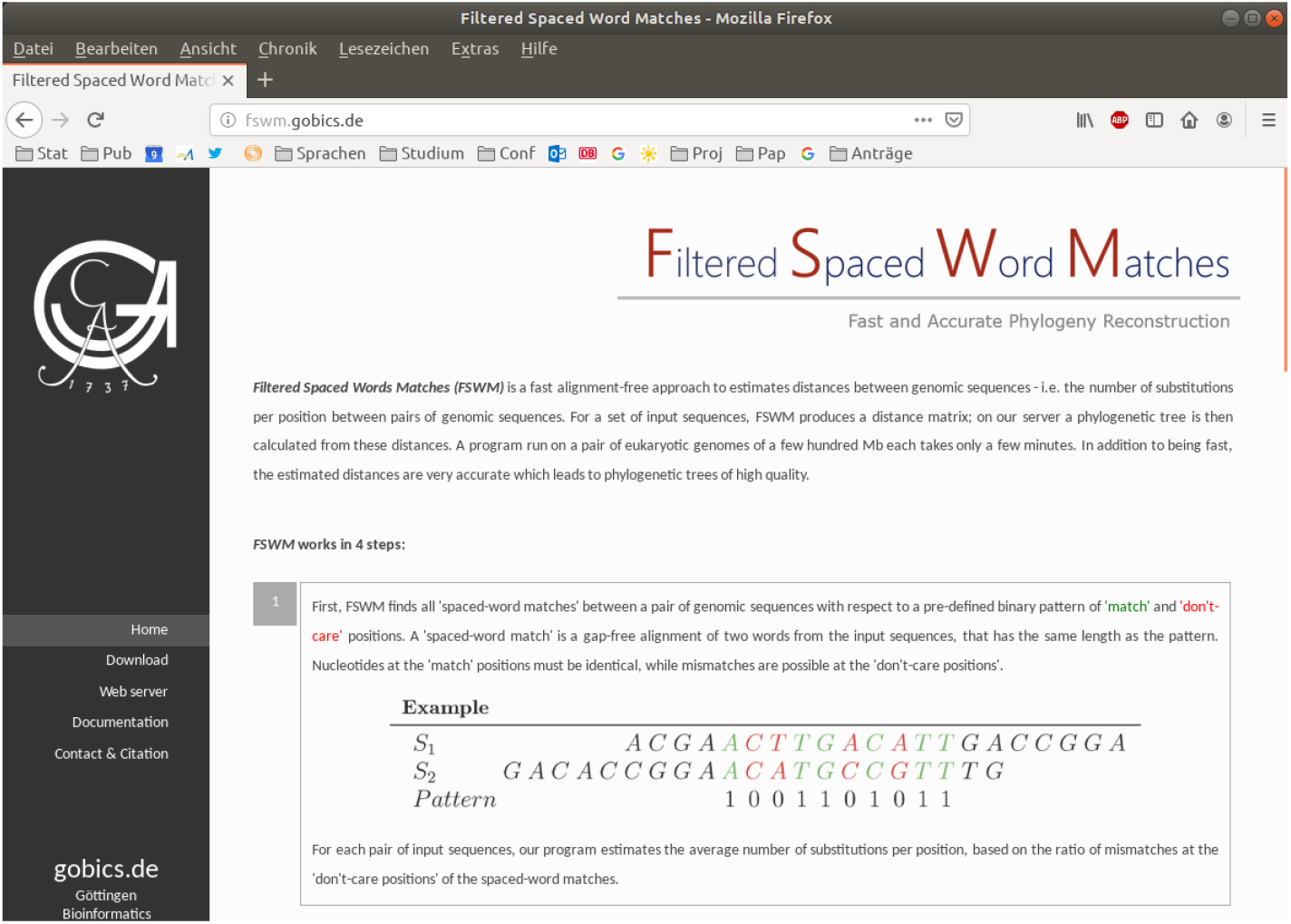
Homepage of *Filtered Spaced Word Matches (FSWM)* at *Göttinen Bioinformatics Compute Server (GOBICS)*

In addition, the above described software tools *FSWM, Prot-SpaM, Multi-SpaM, Read-SpaM* and *Slope-SpaM* are freely available as source code through *github* or through our home page, details are given in Table 1.

**Table 1:** Our software is available as open source code from the following URLs:

|  |  |  |
| --- | --- | --- |
| *Spaced Words* | [55] | http://spaced.gobics.de/ |
| *FSWM* | [47] | http://fswm.gobics.de/ |
| *Prot-SpaM* | [46] | https://github.com/jschellh/ProtSpaM |
| *Multi-SpaM* | [18] | https://github.com/tdencker/multi-SpaM |
| *Read-SpaM* | [42] | https://github.com/burkhard-morgenstern/Read-SpaM |
| *Slope-SpaM* | [66] | https://github.com/burkhard-morgenstern/Slope-SpaM |

## References

[1] Nathan A. Ahlgren, Jie Ren, Yang Young Lu, Jed A. Fuhrman, and Fengzhu Sun. Alignment-free *d*_2_* oligonucleotide frequency dissimilarity measure improves prediction of hosts from metagenomically-derived viral sequences. Nucleic Acids Research, 45:39–53, 2017.

[2] Stephen F. Altschul. Gap costs for multiple sequence alignment. J. Theor. Biol., 138:297–309, 1989.

[3] Samuel V. Angiuoli and Steven L. Salzberg. Mugsy: fast multiple alignment of closely related whole genomes. Bioinformatics, 27:334–342, 2011.

[4] Daniel N. Baker and Ben Langmead. Dashing: fast and accurate genomic distances with HyperLogLog. Genome Biology, 20:265, 2019.

[5] Guillaume Bernard, Cheong Xin Chan, Yao-Ban Chan, Xin-Yi Chua, Yingnan Cong, James M. Hogan, Stefan R. Maetschke, and Mark A. Ragan. Alignment-free inference of hierarchical and reticulate phylogenomic relationships. Briefings in Bioinformatics, 22:426–435, 2019.

[6] B. Edwin Blaisdell. A measure of the similarity of sets of sequences not requiring sequence alignment. Proceedings of the National Academy of Sciences of the United States of America, 83:5155–5159, 1986.

[7] B Edwin Blaisdell. Average values of a dissimilarity measure not requiring sequence alignment are twice the averages of conventional mismatch counts requiring sequence alignment for a computer-generated model system. J Mol Evol, 29:538–47, 1989.

[8] Marcus Boden, Martin Schöneich, Sebastian Horwege, Sebastian Lindner, Chris-André Leimeister, and Burkhard Morgenstern. Alignment-free sequence comparison with spaced *k*-mers. In Tim Beißbarth, Martin Kollmar, Andreas Leha, Burkhard Morgenstern, Anne-Kathrin Schultz, Stephan Waack, and Edgar Wingender, editors, German Conference on Bioinformatics 2013, volume 34 of OpenAccess Series in Informatics (OASIcs), pages 24–34, Dagstuhl, Germany, 2013. Schloss Dagstuhl–Leibniz-Zentrum fuer Informatik.

[9] Karel Břinda, Alanna Callendrello, Lauren Cowley, Themoula Charalampous, Robyn S Lee, Derek R MacFadden, Gregory Kucherov, Justin O’Grady, Michael Baym, and William P Hanage. Lineage calling can identify antibiotic resistant clones within minutes. bioRxiv, 10.1101/403204, 2018.

[10] A. Broder. On the resemblance and containment of documents. In Proceedings of the Compression and Complexity of Sequences 1997, SEQUENCES’97, pages 21–, Washington, DC, USA, 1997. IEEE Computer Society.

[11] Francesca Chiaromonte, Von Bing Yap, and Webb Miller. Scoring pair-wise genomic sequence alignments. In Russ B. Altman, A. Keith Dunker, Lawrence Hunter, and Teri E. Klein, editors, Pacific Symposium on Biocomputing, pages 115–126, Lihue, Hawaii, 2002.

[12] JaeJin Choi and Sung-Hou Kim. Genome tree of life: Deep burst of organism diversity. bioRxiv, 10.1101/756155, 2019.

[13] Benny Chor, David Horn, Yaron Levy, Nick Goldman, and Tim Massingham. Genomic DNA *k*-mer spectra: models and modalities. Genome Biology, 10:R108, 2009.

[14] Matteo Comin and Davide Verzotto. Alignment-free phylogeny of whole genomes using underlying subwords. Algorithms for Molecular Biology, 7:34, 2012.

[15] Aaron C E Darling, Bob Mau, Frederick R Blattner, and Nicole T Perna. Mauve: multiple alignment of conserved genomic sequence with rearrangements. Genome Research, 14:1394–1403, 2004.

[16] Aaron E. Darling, Bob Mau, and Nicole T. Perna. progressiveMauve: Multiple Genome Alignment with Gene Gain, Loss and Rearrangement. PLOS ONE, 5:e11147+, 2010.

[17] Margaret O. Dayhoff, Robert M. Schwartz, and Bruce C. Orcutt. A model of evolutionary change in proteins. Atlas of Protein Sequence and Structure, 6:345–362, 1978.

[18] Thomas Dencker, Chris-André Leimeister, Michael Gerth, Christoph Bleidorn, Sagi Snir, and Burkhard Morgenstern. *Multi-SpaM*: a Maximum-Likelihood approach to phylogeny reconstruction using multiple spaced-word matches and quartet trees. NAR Genomics and Bioinformatics, 2:lqz013, 2020.

[19] Dee R. Denver, Amanda M. V. Brown, Dana K. Howe, Amy B. Peetz, and Inga A. Zasada. Genome Skimming: A rapid approach to gaining diverse biological insights into multicellular pathogens. PLOS Pathogens, 12(8):e1005713, 2016.

[20] Ruud H. Deurenberg, Erik Bathoorn, Monika A. Chlebowicz, Natacha Couto, Mithila Ferdous, Silvia García-Cobos, Anna M.D. Kooistra-Smid, Erwin C. Raangs, Sigrid Rosema, Alida C.M. Veloo, Kai Zhou, Alexander W. Friedrich, and John W.A. Rossen. Application of next generation sequencing in clinical microbiology and infection prevention. Journal of Biotechnology, 243:16 – 24, 2017.

[21] Steven Dodsworth. Genome skimming for next-generation biodiversity analysis. Trends in Plant Science, 20:525 – 527, 2015.

[22] Richard Durbin, Sean R. Eddy, Anders Krogh, and Graeme Mitchison. Biological sequence analysis. Cambridge University Press, Cambridge, UK, 1998.

[23] Robert C. Edgar. MUSCLE: a multiple sequence alignment method with reduced time and space complexity. BMC Bioinformatics, 5:113, 2004.

[24] Olivier Gascuel. BIONJ: an improved version of the NJ algorithm ased on a simple model of sequence data. Molecular biology and evolution, 14:685–695, 1997.

[25] Dan Gusfield. Algorithms on Strings, Trees, and Sequences: Computer Science and Computational Biology. Cambridge University Press, Cambridge, UK, 1997.

[26] Lars Hahn, Chris-André Leimeister, Rachid Ounit, Stefano Lonardi, and Burkhard Morgenstern. *rasbhari*: optimizing spaced seeds for database searching, read mapping and alignment-free sequence comparison. PLOS Computational Biology, 12(10):e1005107, 2016.

[27] Bernhard Haubold. Alignment-free phylogenetics and population genetics. Briefings in Bioinformatics, 15:407–418, 2014.

[28] Bernhard Haubold, Fabian Klötzl, and Peter Pfaffelhuber. andi: Fast and accurate estimation of evolutionary distances between closely related genomes. Bioinformatics, 31:1169–1175, 2015.

[29] Bernhard Haubold, Peter Pfaffelhuber, Mirjana Domazet-Loso, and Thomas Wiehe. Estimating mutation distances from unaligned genomes. Journal of Computational Biology, 16:1487–1500, 2009.

[30] Steven Henikoff and Jora G. Henikoff. Amino acid substitution matrices from protein blocks. Proc. Natl. Acad. Sci. USA, 89:10915–10919, 1992.

[31] Michael Höhl, Stefan Kurtz, and Enno Ohlebusch. Efficient multiple genome alignment. Bioinformatics, 18:312S–320S, 2002.

[32] Michael Höhl, Isidore Rigoutsos, and Mark A. Ragan. Pattern-based phylogenetic distance estimation and tree reconstruction. Evolutionary Bioinformatics Online, 2:359–375, 2006.

[33] Sebastian Horwege, Sebastian Lindner, Marcus Boden, Klaus Hatje, Martin Kollmar, Chris-André Leimeister, and Burkhard Morgenstern. *Spaced words* and *kmacs*: fast alignment-free sequence comparison based on inexact word matches. Nucleic Acids Research, 42:W7–W11, 2014.

[34] Weichun Huang, David M. Umbach, and Leping Li. Accurate anchoring alignment of divergent sequences. Bioinformatics, 22:29–34, 2006.

[35] Lucian Ilie, Silvana Ilie, and Anahita M. Bigvand. SpEED: fast computation of sensitive spaced seeds. Bioinformatics, 27:2433–2434, 2011.

[36] Paul Jaccard. Étude comparative de la distribution florale dans une portion des alpes et des jura. Bulletin del la Société Vaudoise des Sciences Naturelles, 37:547–579, 1901.

[37] Thomas H. Jukes and Charles R. Cantor. Evolution of Protein Molecules. Academy Press, New York, 1969.

[38] Motoo Kimura. The Neutral Theory of Molecular Evolution. Cambridge University Press, 1983.

[39] Fabian Klötzl and Bernhard Haubold. Phylonium: Fast Estimation of Evolutionary Distances from Large Samples of Similar Genomes. Bioinformatics, in press, DOI 10.1093/bioinformatics/btz903.

[40] Gregory Kucherov. Evolution of biosequence search algorithms: a brief survey. Bioinformatics, 35:3547–3552, 2019.

[41] Stefan Kurtz, Adam Phillippy, Arthur L. Delcher, Michael Smoot, Martin Shumway, Corina Antonescu, and Steven L. Salzberg. Versatile and open software for comparing large genomes. Genome Biology, 5:R12+, 2004.

[42] Anna Katharina Lau, Svenja Dörrer, Chris-André Leimeister, Christoph Bleidorn, and Burkhard Morgenstern. *Read-SpaM*: assembly-free and alignment-free comparison of bacterial genomes with low sequencing coverage. BMC Bioinformatics, 20, in press, 2019.

[43] Chris-André Leimeister, Marcus Boden, Sebastian Horwege, Sebastian Lindner, and Burkhard Morgenstern. Fast alignment-free sequence comparison using spaced-word frequencies. Bioinformatics, 30:1991–1999, 2014.

[44] Chris-Andr/’e Leimeister, Thomas Dencker, and Burkhard Morgenstern. Accurate multiple alignment of distantly related genome sequences using filtered spaced word matches as anchor points. Bioinformatics, 35:211–218, 2019.

[45] Chris-André Leimeister and Burkhard Morgenstern. *kmacs*: the *k*-mismatch average common substring approach to alignment-free sequence comparison. Bioinformatics, 30:2000–2008, 2014.

[46] Chris-Andre Leimeister, Jendrik Schellhorn, Svenja Dörrer, Michael Gerth, Christoph Bleidorn, and Burkhard Morgenstern. Prot-SpaM: Fast alignment-free phylogeny reconstruction based on whole-proteome sequences. GigaScience, 8:giy148, 2019.

[47] Chris-André Leimeister, Salma Sohrabi-Jahromi, and Burkhard Morgenstern. Fast and accurate phylogeny reconstruction using filtered spaced-word matches. Bioinformatics, 33:971–979, 2017.

[48] Ming Li, Bin Ma, Derek Kisman, and John Tromp. PatternHunter II: Highly sensitive and fast homology search. Genome Informatics, 14:164–175, 2003.

[49] Ming Li, Bin Ma, Derek Kisman, and John Tromp. PatternHunter II: highly sensitive and fast homology search. Journal of Bioinformatics and Computational Biology, 02:417–439, 2004.

[50] B. Linard, P. Arribas, C. Andújar, A. Crampton-Platt, and A. P. Vogler. Lessons from genome skimming of arthropod-preserving ethanol. Molecular Ecology Resources, 16:1365–1377, 2016.

[51] Burkhard Morgenstern. A space-efficient algorithm for aligning large genomic sequences. Bioinformatics, 16:948–949, 2000.

[52] Burkhard Morgenstern, Oliver Rinner, Saïd Abdeddaïm, Dirk Haase, Klaus Mayer, Andreas Dress, and Hans-Werner Mewes. Exon discovery by genomic sequence alignment. Bioinformatics, 18:777–787, 2002.

[53] Burkhard Morgenstern, Svenja Schöbel, and Chris-André Leimeister. Phylogeny reconstruction based on the length distribution of *k*-mismatch common substrings. Algorithms for Molecular Biology, 12:27, 2017.

[54] Burkhard Morgenstern, Nadine Werner, Sonja J. Prohaska, Rasmus Steinkamp Isabelle Schneider, Amarendran R. Subramanian, Peter F. Stadler, and Jan Weyer-Menkhoff. Multiple sequence alignment with userdefined constraints at GOBICS. Bioinformatics, 21:1271 – 1273, 2005.

[55] Burkhard Morgenstern, Bingyao Zhu, Sebastian Horwege, and Chris-André Leimeister. Estimating evolutionary distances between genomic sequences from spaced-word matches. Algorithms for Molecular Biology, 10:5, 2015.

[56] Saul B. Needleman and Christian D. Wunsch. A general method applicable to the search for similarities in the amino acid sequence of two proteins. J. Mol. Biol., 48:443–453, 1970.

[57] Brian D. Ondov, Gabriel J. Starrett, Anna Sappington, Aleksandra Kostic, Sergey Koren, Christopher B. Buck, and Adam M. Phillippy. Mash Screen: high-throughput sequence containment estimation for genome discovery. Genome Biology, 20:232, 2019.

[58] Brian D Ondov, Todd J Treangen, Páll Melsted, Adam B Mallonee, Nicholas H Bergman, Sergey Koren, and Adam M Phillippy. Mash: fast genome and metagenome distance estimation using minhash. Genome Biology, 17:132, 2016.

[59] Benedict Paten, Dent Earl, Ngan Nguyen, Mark Diekhans, Daniel Zerbino, and David Haussler. Cactus: Algorithms for genome multiple sequence alignment. Genome Research, 21:1512–1528, 2011.

[60] Enrico Petrucci, Laurent Noé, Cinzia Pizzi, and Matteo Comin. Iterative spaced seed hashing: Closing the gap between spaced seed hashing and *k*-mer hashing. Journal of Computational Biology, in press, doi:10.1089/cmb.2019.0298.

[61] Cinzia Pizzi. MissMax: alignment-free sequence comparison with mismatches through filtering and heuristics. Algorithms for Molecular Biology, 11:6, 2016.

[62] Gesine Reinert, David Chew, Fengzhu Sun, and Michael S. Waterman. Alignment-free sequence comparison (I): Statistics and power. Journal of Computational Biology, 16:1615–1634, 2009.

[63] Jie Ren, Xin Bai, Yang Young Lu, Kujin Tang, Ying Wang, Gesine Reinert, and Fengzhu Sun. Alignment-free sequence analysis and applications. Annual Review of Biomedical Data Science, 1:93–114, 2018.

[64] Sandy Richter, Francine Schwarz, Lars Hering, Markus Böggemann, and Christoph Bleidorn. The utility of genome skimming for phylogenomic analyses as demonstrated for glycerid relationships (Annelida, Glyceridae). Genome Biology and Evolution, 7:3443–3462, 2015.

[65] David F Robinson and Les Foulds. Comparison of phylogenetic trees. Mathematical Biosciences, 53:131–147, 1981.

[66] Sophie Röhling, Alexander Linne, Jendrik Schellhorn, Morteza Hosseini, Thomas Dencker, and Burkhard Morgenstern. The number of *k*-mer matches between two DNA sequences as a function of *k* and applications to estimate phylogenetic distances. bioRxiv, doi:10.1101/527515v3, 2019.

[67] Naruya Saitou and Masatoshi Nei. The neighbor-joining method: a new method for reconstructing phylogenetic trees. Molecular Biology and Evolution, 4:406–425, 1987.

[68] Shahab Sarmashghi, Kristine Bohmann, M. Thomas P. Gilbert, Vineet Bafna, and Siavash Mirarab. Skmer: assembly-free and alignment-free sample identification using genome skims. Genome Biology, 20:34, 2019.

[69] Fabian Sievers, Andreas Wilm, David Dineen, Toby J Gibson, Kevin Karplus, Weizhong Li, Rodrigo Lopez, Hamish McWilliam, Michael Remmert, Johannes Söding, Julie D Thompson, and Desmond G Higgins. Fast, scalable generation of high-quality protein multiple sequence alignments using Clustal Omega. Molecular Systems Biology, 7:539, 2011.

[70] Gregory E. Sims, Se-Ran Jun, Guohong A. Wu, and Sung-Hou Kim. Alignment-free genome comparison with feature frequency profiles (FFP) and optimal resolutions. Proceedings of the National Academy of Sciences, 106:2677–2682, 2009.

[71] Sagi Snir and Satish Rao. Quartet MaxCut: A fast algorithm for amalgamating quartet trees. Molecular Phylogenetics and Evolution, 62:1 – 8, 2012.

[72] Kai Song, Jie Ren, Gesine Reinert, Minghua Deng, Michael S. Waterman, and Fengzhu Sun. New developments of alignment-free sequence comparison: measures, statistics and next-generation sequencing. Briefings in Bioinformatics, 15:343–353, 2014.

[73] Kai Song, Jie Ren, Zhiyuan Zhai, Xuemei Liu, Minghua Deng, and Fengzhu Sun. Alignment-free sequence comparison based on next-generation sequencing reads. Journal of Computational Biology, 20:64–79, 2013.

[74] Alexandros Stamatakis. RAxML version 8: a tool for phylogenetic analysis and post-analysis of large phylogenies. Bioinformatics, 30:1312–1313, 2014.

[75] Hanno Teeling, Anke Meyerdierks, Margarete Bauer, Rudolf Amann, and Frank O. Glöckner. Application of tetranucleotide frequencies for the assignment of genomic fragments. Environmental Microbiology, 6:938–947, 2004.

[76] Sharma V. Thankachan, Sriram P. Chockalingam, Yongchao Liu, and Ambujam Krishnan Srinivas Aluru. A greedy alignment-free distance estimator for phylogenetic inference. BMC Bioinformatics, 18:238, 2017.

[77] Igor Ulitsky, David Burstein, Tamir Tuller, and Benny Chor. The average common substring approach to phylogenomic reconstruction. Journal of Computational Biology, 13:336–350, 2006.

[78] Susana Vinga and Jonas Almeida. Alignment-free sequence comparison - a review. Bioinformatics, 19:513–523, 2003.

[79] Susana Vinga, Alexandra M. Carvalho, Alexandre P. Francisco, Luís M. S. Russo, and Jonas S. Almeida. Pattern matching through Chaos Game Representation: bridging numerical and discrete data structures for biological sequence analysis. Algorithms for Molecular Biology, 7:10, 2012.

[80] Lin Wan, Gesine Reinert, Fengzhu Sun, and Michael S Waterman. Alignment-free sequence comparison (II): theoretical power of comparison statistics. Journal of Computational Biology, 17:1467–1490, 2010.

[81] Kevin Weitemier, Shannon C. K. Straub, Richard C. Cronn, Mark Fishbein, Roswitha Schmickl, Angela McDonnell, and Aaron Liston. Hyb-seq: Combining target enrichment and genome skimming for plant phylogenomics. Applications in Plant Sciences, 2:1400042, 2014.

[82] Huiguang Yi and Li Jin. Co-phylog: an assembly-free phylogenomic approach for closely related organisms. Nucleic Acids Research, 41:e75, 2013.

[83] Andrzej Zielezinski, Hani Z Girgis, Guillaume Bernard, Chris-Andre Leimeister, Kujin Tang, Thomas Dencker, Anna Katharina Lau, Sophie Röhling, JaeJin Choi, Michael S Waterman, Matteo Comin, Sung-Hou Kim, Susana Vinga, Jonas S Almeida, Cheong Xin Chan, Benjamin James, Fengzhu Sun, Burkhard Morgenstern, and Wojciech M Karlowski. Bench-marking of alignment-free sequence comparison methods. Genome Biology, 20:144, 2019.

[84] Andrzej Zielezinski, Susana Vinga, Jonas Almeida, and Wojciech M. Karlowski. Alignment-free sequence comparison: benefits, applications, and tools. Genome Biology, 18:186, 2017.

